# Structural basis of the transcription termination factor Rho engagement with transcribing RNA polymerase

**DOI:** 10.1101/2022.09.02.506315

**Authors:** Yuko Murayama, Haruhiko Ehara, Mari Aoki, Mie Goto, Takeshi Yokoyama, Shun-ichi Sekine

## Abstract

Transcription termination is an essential step in transcription by RNA polymerase (RNAP) and crucial for gene regulation. For many bacterial genes, transcription termination is mediated by the ATP-dependent RNA translocase/helicase Rho, which causes the dissociation of RNA/DNA from RNAP elongation complex (EC). However, structural basis of the interplay between Rho and RNAP remains obscure. Here we report the cryo-electron microscopy structure of the Rho-engaged EC. The Rho hexamer binds RNAP through the C-terminal domains, which surround the RNA-exit site of RNAP, directing the nascent RNA seamlessly from the RNA exit to the Rho central channel. The β-flap tip at the RNA exit is critical to the Rho-dependent RNA release, and its deletion causes an alternative Rho-RNAP binding mode, which is irrelevant to termination. The Rho-binding site overlaps with the binding sites of other macromolecules, such as ribosomes, providing a general basis of gene regulation.

**Teaser:** Cryo-EM captures the structure of an RNA polymerase elongation complex engaged with the termination factor Rho.

## Introduction

From bacteria to eukaryotes, DNA-dependent RNA polymerase (RNAP) conducts gene transcription via three sequential processes, initiation, elongation, and termination. The last termination process includes stalling of transcribing RNAP elongation complex (EC) at termination sites within a gene, followed by dissociation of RNA and DNA from RNAP to generate the proper RNA 3’ end. Two types of mechanisms are known for termination: intrinsic termination that does not require any auxiliary factors and factor-dependent termination that requires a specific protein(s) for termination(*1*). In bacteria, the factor-dependent termination is mediated by transcription termination factor Rho, which terminates transcription at Rho-dependent terminators of many genes(*2–5*). Rho plays a crucial role in defining the end of the genes and gene boundaries. Besides its basic role in the RNA synthesis, emerging evidences have revealed implication of Rho in key regulatory mechanisms, including transcription suppression of untranslated mRNA (the polarity effects)(*6*) and small RNA or riboswitch-dependent regulation(*7, 8*). Rho-dependent termination is also implicated in “housekeeping” mechanisms, such as suppression of pervasive antisense transcription(*9*), silencing of horizontally transferred genes(*10*), and resolving conflicts between transcription and replication(*11*).

Rho is an ATP-dependent RNA translocase/helicase with a 5’→3’ directionality(*12, 13*). It consists of the N-terminal domain (NTD) and the C-terminal domain (CTD), which possess the primary RNA binding site (PBS) and the secondary RNA binding site (SBS), respectively(*14–16*). Rho forms a ring-shaped homo-hexameric complex mediated by CTD(*16, 17*). The Rho hexamer can take either open- or closed-ring conformation. When Rho binds RNA and ATP, it assumes the closed-ring conformation to thread the RNA into its central channel composed of SBS(*18*). At the Rho-dependent terminator, Rho engages nascent RNA that is emerging from EC, and causes dissociation of RNA and DNA from RNAP(*19–21*). However, the mechanism by which Rho interacts with RNAP to cause termination remains elusive.

Recently, cryo-electron microscopy (cryo-EM) structures of *Escherichia coli* RNAP bound with Rho and transcription elongation factor NusA and/or NusG were reported(*22, 23*). In these structures, the Rho hexamer was in an open conformation without engagement with RNA, and binds RNAP via an *E. coli* specific insertion domain (βSI2) and NusA. These structures suggested that Rho induces an allosteric change in RNAP to destabilize EC for priming the dissociation of RNA and DNA. However, these studies also postulated that termination ultimately involves the Rho ring closure and its engagement with RNA, followed by the Rho translocation to release RNA and DNA. In this study, we recapitulated the Rho-associated EC by using proteins from the bacterium *Thermus thermophilus*, and performed structure determination by cryo-EM single particle analysis. The structure revealed the manner by which the nascent RNA-engaged Rho hexamer in a closed-ring conformation interacts with the RNA-exit site of RNAP, illuminating the EC state prior to termination.

## Results

### Rho-dependent RNA-release assay

To examine the Rho-dependent RNA release activity from transcription elongation complex (EC) of RNAP, we developed an *in vitro* experimental system by using *Thermus thermophilus* Rho and RNAP. A 180 bp DNA, which contains a 114 bp G-less segment, was designed, and one end of the DNA was immobilized to a magnetic bead (Fig. 1A, S1A). The other end of the DNA has a 3’ overhang of the template strand, to which a fluorescently-labeled primer RNA was annealed, and RNAP was loaded to assemble EC. A 125-nucleotide RNA was elongated by adding ATP, CTP, and UTP, and EC was stalled at the first G stretch on the DNA. After washing out the nucleotide substrates, Rho was loaded to form an EC-Rho complex. When ATP was added, RNA was released from the immobilized complex to the solvent, indicating the capability of this complex in termination (Fig. 1B). The RNA release is ATP dependent, as it did not occur in the absence of ATP or in the presence of an ATP analog, ADP-aluminum fluoride (ADP•AlF_4_) or ADP-beryllium fluoride (ADP•BeF_3_).

**Fig. 1.**
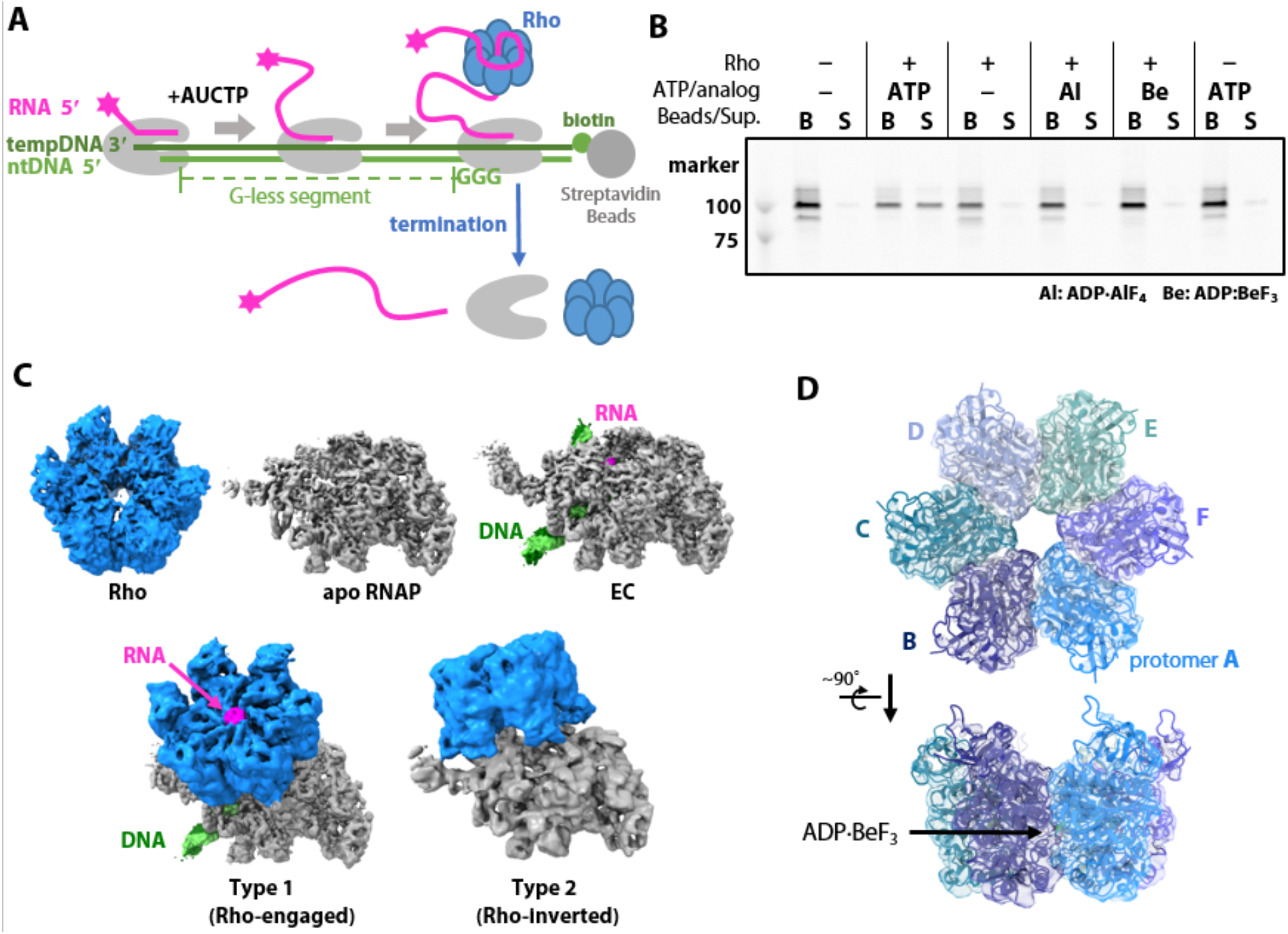
Reconstitution of the Rho-EC complex and its cryo-EM analysis. (A) Schemes of the Rho-EC reconstitution and the RNA release assay. (B) Rho-dependent RNA release assay. Rho and ATP (or ADP•AlF_4_ or ADP•BeF_3_) was added to ECs immobilized on streptavidin magnetic beads, and the mixture was incubated for 10 minutes at 65°C. Each sample was separated into beads (B) and supernatant (S) fractions, and analyzed by urea-PAGE. RNA was detected by 5’-fluorescent label. (C) Structures of the complexes contained in the cryo-EM sample. Cryo-EM maps of Rho, apo RNAP, EC, and type-1 and type-2 complexes are shown. The maps are colored to match the structural models (grey, RNAP; blue, Rho; green, DNA; magenta, RNA). (D) Structure of *T. thermophilus* Rho. The Rho cartoon model is shown in two orientations, with transparent cryo-EM map overlaid.

### Cryo-EM analysis of the RNAP-Rho complexes

For the cryo-EM analysis of the EC-Rho complex, EC was prepared using the same DNA/RNA as in the RNA-release assay. The nucleotide substrates and RNAP-unbound DNA/RNA were removed from EC by gel filtration chromatography (Fig. S1B). The purified EC was mixed with Rho in presence of ADP•BeF_3_, and crosslinked with BS3 to stabilize the complex before preparation of cryogrids. Approximately 33,000 cryo-EM images were collected for single particle analysis (Fig. S1C). By the initial 2D classification, the particles were classified into the Rho-only and RNAP-containing classes (Fig. 1C, S1D, E). The Rho structure was determined at 3.5 Å resolution (Fig.1C, S2). The Rho hexamer assumes a closed-ring conformation with the ATP analog ADP·BeF_3_ bound at the interface between the protomers (Fig. 1D, Fig. S1F). The closed-ring structure is similar to the previously reported crystal structure of the *E. coli* Rho closed hexamer bound with ADP•BeF_3_ and RNA (C_α_ r.m.s.d = 1.9 Å; Fig. S1G) (PDB 5JJI) (*17*).

The particles containing RNAP were further classified by 3D classification into DNA/RNA-unbound RNAP (apo RNAP), EC, and two forms of RNAP-Rho complexes (referred to as the type-1 and type-2 complexes) (Fig. S2, S3). EC holds a transcription bubble containing a 10 base-pair DNA/RNA hybrid within the RNAP main channel (Fig.1C,2). The EC is stalled immediately before the G-triad of the template DNA (Fig. 1A). The type-1 and type-2 RNAP-Rho complexes both reveal the closed-ring Rho hexamer with RNA threaded in the central channel, but they are different in the binding sites and orientations of Rho relative to RNAP. In the type-1 complex, Rho is bound to the RNA exit site of EC via its CTD side, representing the Rho-engaged state of the EC. By contrast, in the type-2 complex, Rho is bound to apo RNAP via its NTD side, probably representing a non-productive state.

**Fig. 2.**
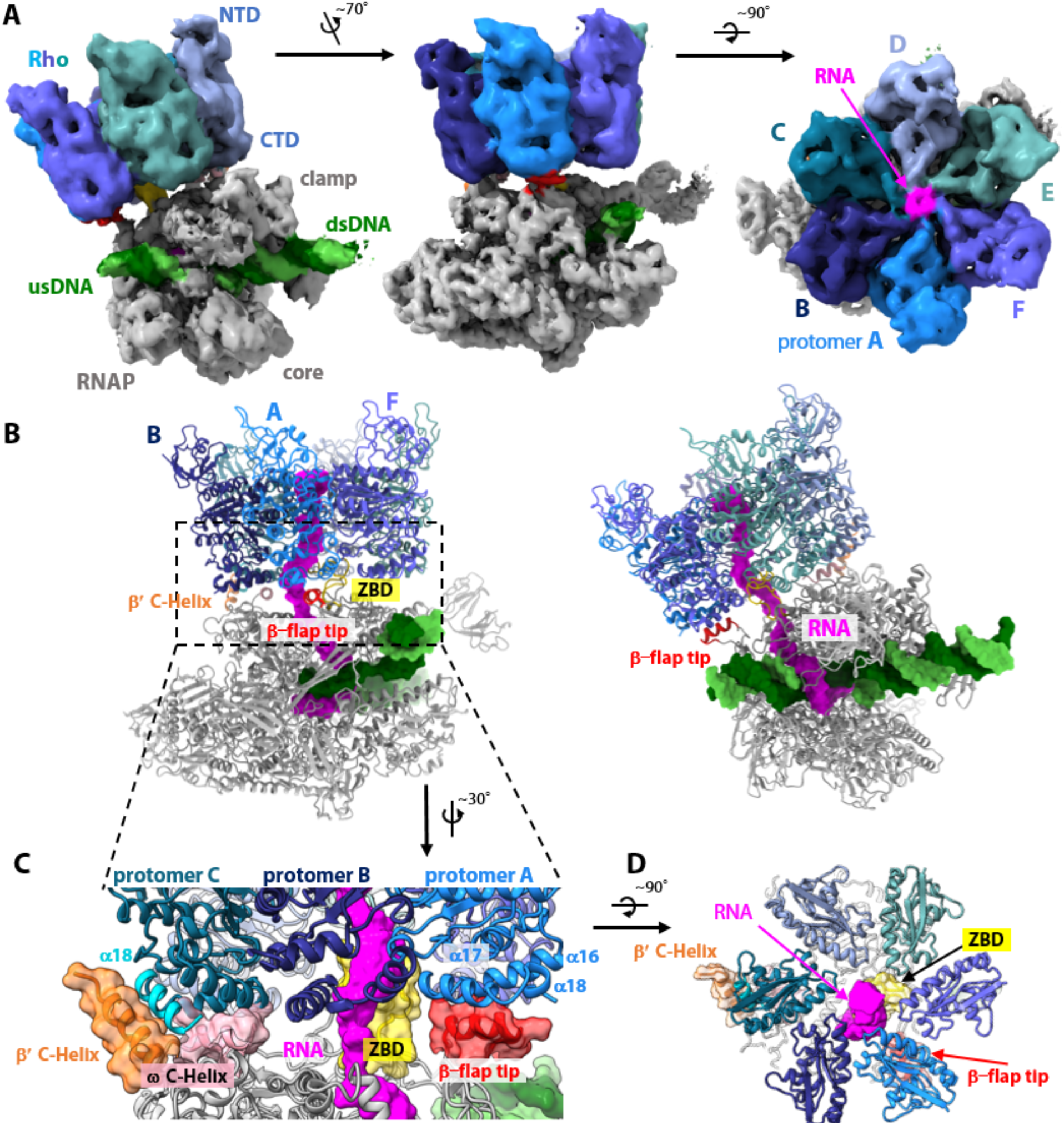
Structure of the Rho-engaged EC (Type 1) (A) Cryo-EM map of the Rho-engaged EC (the type-1 complex) in three orientations. The composite map was generated from the EC-masked (3.7 Å resolution) and Rho-masked (4.6 Å resolution) focused maps by Phenix(*39*). The map is colored to match the model colors; Rho is colored as in Fig. 1D, RNAP is colored gray (β-flap tip, β’ C-helix, and zinc-binding domain (ZBD) in red, orange, and yellow, respectively), template DNA in dark green, non-template DNA in light green, and RNA in magenta. (B) Structure of type-1 complex. The structural model is shown in two orientations, with the protein shown in cartoon representation and the DNA/RNA shown in surface representation. (C) Close-up views of the RNAP-Rho interface. Transparent surface of β-flap tip, ZBD, β’ C-helix and ω C-helix domains are overlaid. (D) The Rho hexamer ring from the NTD side.

### Structure of the Rho-engaged EC (Type 1)

In the type-1 complex, the Rho hexamer assumes a closed-ring conformation and is engaged with EC (Fig. 2). Rho directly binds the RNAP RNA-exit site via its CTD side, so that the Rho ring surrounds the exiting nascent RNA to form a continuous RNA path from the RNAP RNA exit channel to the Rho central channel. The CTD side of the Rho hexamer is widened as compared with that of the free Rho hexamer (Fig. S4A). The nascent RNA emerging from RNAP is seamlessly threaded into the Rho central channel (the secondary binding site (SBS)).

The β-flap domain of the RNAP β subunit (residues β762-784) forms a part of the RNA exit channel. The tip helix of the β-flap domain is reoriented and adapted to the C-terminal side of a Rho protomer (protomer A in Fig. 2), contacting helices α16 (residues 381-395), α17 (residues 399-412), and α18 (residues 415-419). Thereby, the position of the β-flap tip helix is slightly away from the exiting RNA, as compared with its position in the previously reported crystal structure (PDB 2O5I)(*24*) (Fig. S4B). The zinc-binding domain of the β’ subunit (residues β’56-80), which is also a part of the RNA exit channel, enters into the central channel of the Rho ring, like an axle of a wheel. Rho also binds to the C-terminus of the β’ subunit and the ω subunit of RNAP. The C-terminal helix (helix a 18; residues 415-426) of a Rho protomer (protomer C) folds longer than that in the other protomers (residues 415-420), and forms a helix bundle with the C-terminal helix of the RNAP β’ subunit (residues β’1489-1502). This helix bundle formation displaces the C-terminal helix of the RNAP ω subunit (residues ω 82-92), which instead contacts the bottom part of the Rho protomer C (Fig. 2D, S4C). Thus, the nascent RNA, emerging between the zinc-binding and the β-flap domains, is seamlessly directed to the Rho central channel, passing through the narrowest part consisting of SBSs of the Rho protomers (Fig. 2).

### The β-flap tip is critical for the Rho-dependent RNA release and the type-1 complex formation

To investigate the significance of the Rho-contacting RNAP domains in transcription termination, mutant RNAPs lacking the β-flap tip or the zinc-binding domain were created, and their Rho-dependent RNA release activities were examined. While deletion of the zinc-binding domain did not impair the RNA release activity, deletion of the β-flap tip substantially impaired the activity (Fig.3A). Thus, the β-flap tip is crucial for the Rho-dependent RNA release.

**Fig. 3.**
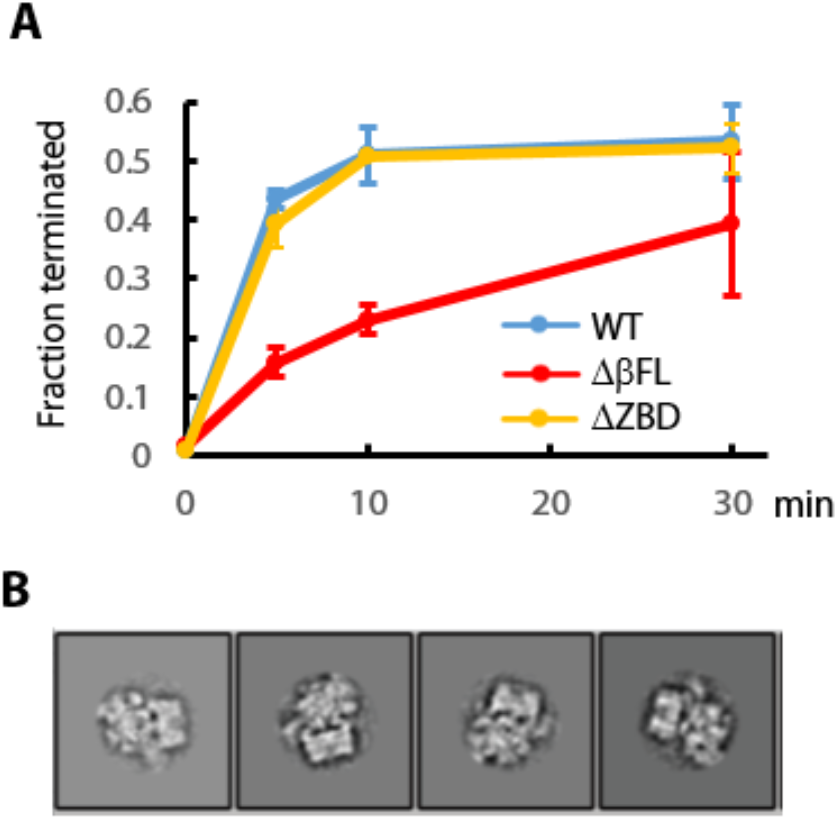
Rho-dependent RNA release assays with RNAP variants. (A) Rho-dependent RNA release activity were examined with RNAP variants. Mean values of three independent experiments were plotted (error bars = S.D.). (B) 2D-class averages of ~3,000 particles of RNAP-Rho complex prepared using Δβ-flap-tip RNAP variant.

We also performed cryo-EM analysis of the EC-Rho complex prepared using the β-flap-tip deleted RNAP. The structure analysis yielded only the type-2 complex class, but no type-1 complex class (Fig. 3B). This suggests that the β-flap tip plays a critical role in the type-1 complex formation through interaction with the C-terminal part of Rho and the type-1 complex is implicated in termination.

### Structure of RNAP bound with an inverted Rho (Type 2)

In the type-2 complex, Rho is bound to RNAP in an inverted orientation, as compared with that in the type-1 complex (Fig 4). The Rho hexamer assumes a closed-ring conformation with RNA threaded through the Rho central channel, and binds near the DNA-exit site of RNAP through its NTD side. The RNAP reveals an open clamp conformation, which resembles that in apo RNAP than EC (Fig S3D). While the RNAP main channel is empty, with no bound DNA/RNA hybrid, the Rho-bound RNA appears to enter and interact with the upstream DNA binding site of the RNAP (Fig. S6). Four out of the six Rho protomers are involved in the RNAP interaction; the Rho protomers B and C contact the base part of the β-flap domain (residues β727-738), the protomers C and D contact the jaw-lobe 1 domain, and the protomer E contacts the clamp coiled-coil of RNAP.

**Fig. 4.**
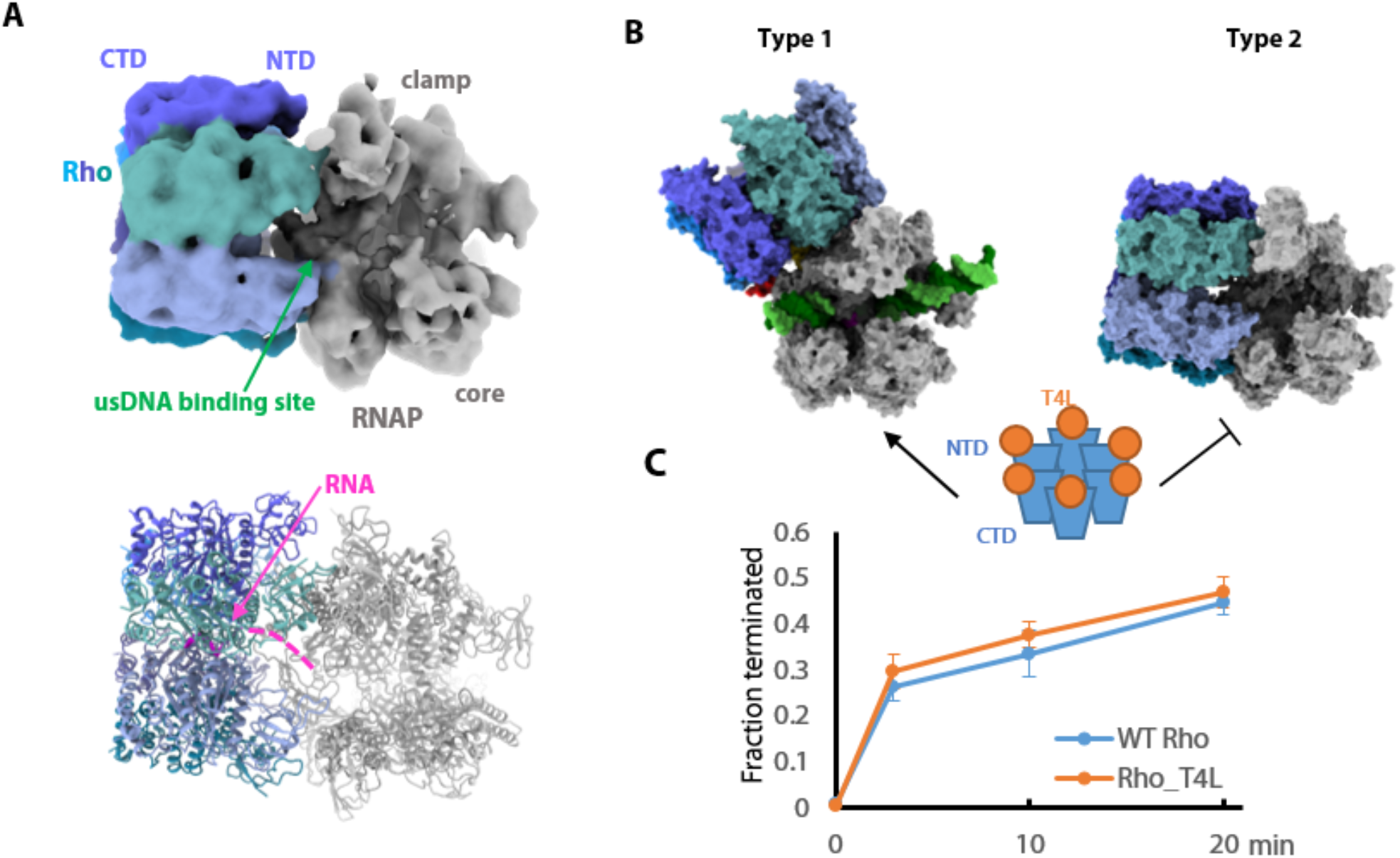
Structure of RNAP bound with an inverted Rho (Type 2) (A) Structure of the type-2 complex. A 5.8 Å cryo-EM map (top) and the structural model (bottom). The map is colored as in Fig. 2. (B) Comparison of the type-1 and type-2 complexes. The surface models of the two complexes are shown with the same RNAP orientation. (C) The RNA release assay with the T4L-inserted Rho. Mean values of three independent experiments were plotted (error bars = S.D.).

To investigate whether the observed inverted binding mode of Rho is significant, we created a mutant Rho in which T4 lysozyme (T4L) is inserted into the Rho NTD (between residues 115 and 116), and examined the RNA-release activity. As the bulky T4L insertion would intervene between the Rho NTD and RNAP, it is expected to hinder the inverse RNAP-Rho interaction as observed in the type-2 complex, but not the RNAP-Rho interaction as observed in the type-1 complex (Fig. 4C). The T4L-inserted Rho mutant showed an RNA release activity comparable to that of the wild-type Rho. This suggests that the inverted binding mode in the type-2 complex is not required for the RNA release. The type-2 complex might not be relevant to termination, but represent another state, such as a post-termination state that is formed by interaction between Rho and apo RNAP after dissociation of DNA/RNA from EC.

## Discussion

In this study, we determined the cryo-EM structure of a Rho-engaged EC, in which the Rho hexamer is in a closed-ring conformation and engaged with RNA and EC (the type-1 complex). The Rho hexamer binds to the rim of the RNA-exit channel of EC via its CTD side, and seamlessly threads the emerging nascent RNA into its central channel. The β-flap tip, which forms part of the RNA exit channel, was shown to be essential for the Rho-EC engagement and the Rho-dependent RNA release. Thus, Rho binding by EC is mediated by both nascent RNA and the specific Rho-RNAP interaction, consistent with the “classic” view of the Rho-dependent transcription termination.

On the other hand, our cryo-EM analysis provided another Rho-RNAP complex (the type-2 complex), in which Rho binds near the DNA exit channel of apo RNAP through its NTD side. The T4L insertion into NTD did not affect the RNA release activity of Rho. Moreover, deletion of the RNAP β-flap tip, which is critical to the Rho-dependent RNA release, enriched the type-2 complex. Therefore, the inverted Rho-binding mode in this complex is unlikely to be relevant to termination. As Rho preferentially binds RNAP in this alternative mode, this complex could represent a state formed after EC has released DNA and RNA.

In the previous studies of *E. coli* Rho-RNAP complex structures, Rho binds near the DNA exit site of EC through its NTD side, as in our inverted Rho-RNAP structure (the type-2 complex) (Fig. S7) (*22, 23*). However, in the *E. coli* complexes, the Rho hexamer assumes an open-ring conformation without full engagement with RNA, and the Rho-RNAP interaction was mediated by an *E. coli* specific insertion (βSI2) of RNAP and NusA. While these studies postulated that the Rho-RNAP interaction induces allosteric change in the EC active site, they also proposed that transcription termination occurs via its subsequent state in that Rho assumes a closed-ring conformation to gain access to both nascent RNA and the RNA-exit channel of RNAP. Our current structure of the Rho-engaged EC (the type-1 complex) might have captured this state.

The Rho-RNAP binding mode in the current Rho-engaged EC structure is also consistent with the NusG-dependent Rho recruitment. The transcription elongation factor NusG recruits Rho to EC, and facilitates transcription termination in *E. coli*(*4*). The Rho CTD becomes the interface with the NusG CTD, and their interaction promotes termination. Although *T. thermophilus* NusG did not bind either EC or Rho, modeling of an *E. coli* EC-Rho complex based on the current type-1 complex structure suggests that the NTD and CTD of NusG could bridge between RNAP and Rho, respectively, reasonably explaining the NusG-mediated recruitment of Rho to the EC (Fig. S8).

How does Rho terminate transcription? If Rho “pulls” RNA by its ATP-dependent RNA translocation activity as suggested earlier, it would invoke perturbation in the DNA/RNA hybrid within the transcription bubble or in RNAP-DNA/RNA contacts (*19*). This would trigger destabilization of EC, leading to the RNAP conformational change including the clamp opening, the bubble collapse, and release of DNA/RNA from the RNAP(*25*). When the apo RNAP structure is superimposed with the Rho-engaged EC structure, the opened clamp in apo RNAP fits to the bottom parts of the Rho hexamer without severe steric clash (protomers E and F). Therefore, the Rho-engaged EC is compatible with the clamp opening of RNAP, for the release of the bound DNA/RNA (Fig. S9).

In this study, Rho was revealed to bind directly to the RNA-exit site of RNAP. In bacteria, RNAP interacts with the trailing ribosome to establish the transcription-translation coupling(*26–29*). Once the ribosome reaches the stop codon and detaches from the preceding RNAP, Rho can access the RNAP to terminate transcription. The ribosome-binding site of RNAP overlaps with the Rho-binding site, suggesting that their RNAP bindings are mutually exclusive (Fig. S10). This reasonably explains the coordination of transcription and translation in gene regulation. The Rho-binding site is also overlapped with that for the antitermination complex(*30, 31*), explaining how the latter blocks the Rho access to the RNAP transcribing ribosomal genes or phage operons. Thus, the current structure provides structural foundations of gene regulation via coordination or competition of multiple molecular processes.

## Materials and Methods

### Proteins

For the expression of recombinant *T. thermophilus* RNAP, the genes of the RNAP subunits were cloned into pETDuet-1 (β and β’ subunits in MCS1, a subunit in MCS2; a FLAG tag was added to the C-terminal end of the β’ subunit) and pET-47b (ω subunit) vectors. These plasmids were co-expressed in *Escherichia coli* Rosetta2 (DE3). Plasmids for expression of RNAP variants were constructed based on these plasmids. In Δβ-flap-tip and ΔZBD, the β-flap tip helix (residues β762-782) and the zinc binding domain (residues β’57-80), respectively, were removed, and the gaps were filled with three alanine residues. The recombinant RNAP variants were purified as described previously(*32*).

*T. thermophilus* Rho gene was cloned into pET-47b vector. The N-terminally His-tagged Rho was expressed in *E. coli* Rosetta2 (DE3). The cells were suspended in the lysis buffer A (50 mM Hepes-NaOH pH 7.5, 500 mM KCl, 10% glycerol, 1 mM DTT), and disrupted by sonication. The lysate was heated at 70°C for 10 minutes to denature the *E. coli* proteins, and the precipitate was removed by centrifugation. The supernatant was loaded onto a Ni affinity column (Ni-Sepharose 6 FF, Invitrogen). The column was washed with the lysis buffer A supplemented with 10 mM imidazole. The His-tag was cleaved by the HRV 3C protease on the column for 16 hours at 4°C. The flow through fraction after the HRV 3C cleavage was purified by a gel filtration column (Superdex 200, Cytiva) equilibrated with the buffer A.

The C-terminally His-tagged Rho (Rho_CHis) was cloned into pET-22b vector. Protein expression, cell lysis and heat treatment were performed as for the N-terminally His-tagged Rho. The supernatant of the heat treatment was loaded onto a Ni affinity column, and the Rho_CHis protein was eluted with lysis buffer A supplemented with 500 mM imidazole. The Rho_CHis protein was precipitated by lowering the NaCl concentration from 500 mM to 100 mM. The precipitate was re-dissolved in lysis buffer A, and purified by gel filtration column chromatography (Superose 6, Cytiva). Rho_CHis was used for the RNA release assay with RNAP variants (Fig. 3A). In other assays (Figs. 1B, 4C) and structural analysis, N-terminal His-tag-removed Rho was used.

The Rho variant with a T4 lysozyme insertion (Rho_T4L) was constructed by inserting the T4 lysozyme gene between residues Glu115 and Asn116 of the Rho gene, and was cloned into pET-47b vector. The Rho_T4L protein was expressed and purified similarly to the N-terminally His-tagged Rho, except that the lysate was not heat-treated.

*T. thermophilus* GreA gene was cloned into pET-22b vector. The C-terminally His-tagged GreA was expressed in *E. coli* Rosetta2 (DE3). The cells were disrupted in lysis buffer B (50 mM Tris-HCl pH 8.0, 150 mM NaCl, 1 mM DTT). The lysate was heated at 80°C for 10 minutes, and the supernatant was loaded onto a Ni affinity column (Ni-sepharose 6 FF, Invitrogen). GreA was eluted with the lysis buffer supplemented with 500 mM imidazole, and was dialyzed against the lysis buffer B.

*T. thermophilus* NusG gene was cloned into pET-47b vector. The N-terminally His-tagged NusG was expressed in *E. coli* Rosetta2 (DE3). The cells were disrupted in the lysis buffer B. The lysate was heated at 80°C for 10 minutes, and the supernatant was loaded onto a Ni affinity column (Ni-Sepharose 6 FF, Invitrogen). The His-tag was cleaved by the HRV3C protease on the column, and NusG was eluted with the lysis buffer B.

### Rho-dependent termination assay

The elongation complex of RNAP was prepared by walking the RNAP on a template DNA (Fig. 1A, S1A). The template DNA sequence was amplified using primers T1 (CCACAGCCAGGATCCTGCAGGACTTCACTCCCTACTCAACTACTATCTACCCATCTCTCTTCACTCCATA), T2 (GAGATATGGTTATGGGTAAGATAGATGGTGAGGTGATG AGTTTAAAGGAGTGAAGTATGGAGTGAAGAGA), T3 (CCATAACCATATCTCCACAT CCACCTGGGTGCTTGTGGTAGTGCACCGATCCCTGTCGACTTTCCCGT), and T4 (TTTACCAGACTCGAGACGGGAAAGTCGACAGG). The resulting ~200mer DNA was cloned between the BamHI and XhoI sites of the pETDuet-1 vector. Biotinylated template DNA was amplified using primers T0-Fw (CCACAGCCAGGATCCTGCAGGACTTCACTCC) and T3-Rv-bio (GAACGCATTACCAGAGAATTCACGGGAAAGTCGACAGGGATC, 5’-biotinylated). About 1 pmol per sample of the biotinylated template DNA was captured on the streptavidin magnetic beads (Dinabeads MyOne streptavidin C1, Invitrogen) in buffer A (40 mM Tris-HCl pH 7.4, 1 M NaCl). The DNA was digested by PstI for 1 hour at 37°C in the H buffer (50 mM Tris-HCl pH 7.5, 100 mM NaCl, 10 mM MgCl_2_, 1 mM DTT) to make a 3’-protruding end in the upstream DNA end. After digestion, the beads were washed with the M buffer (50 mM Tris-HCl pH 7.5, 50 mM NaCl, 10 mM MgCl_2_, 1 mM DTT). For the elongation complex (EC) reconstitution, RNAP, fluorescently labeled primer RNA (AAUUUGCAGGA; 5’-DY647 labeled), NusG, GreA, and substrates ATP/CTP/UTP were added to the beads, and the mixture was incubated for 15 minutes at 65°C. The molar ratio of the template DNA: RNAP: primer: NusG: GreA was approximately 1:2:2:4:4. The beads were washed for two times by the H buffer, and the remaining EC was pre-incubated for one minute at 35°C or 65°C for the subsequent Rho-dependent termination assay. For termination, a mixture of Rho (1 pmol/sample) and ATP (final 1 mM) were added to the EC on the beads. 5 μl aliquots were withdrawn at the indicated time points, and the supernatant was separated from the beads by standing the tubes for 1 minute on a magnetic stand. 5 μl of the urea-PAGE sample buffer (50 mM Tris-HCl pH 7.5, 50 mM EDTA, 5 M urea, 1% orange G) was mixed with the sample, and the RNAs were fractionated by 20% PAGE containing 7 M urea. The fluorescent-labeled RNA was imaged and quantitated with ImageQuant LAS-4000 (GE Healthcare) or Typhoon imager (Amersham). The same reaction was performed in the absence of ATP or in the presence of ATP analogs. The ATP mimic ADP·AlF_4_ was prepared by mixing ADP, AlCl_3_, and NaF in a molar ratio of 1:5:20. The ATP mimic ADP•BeF_3_ was prepared by mixing ADP, BeSO_4_ and NaF in a molar ratio of 1:5:15.

### EC-Rho preparation for cryo-EM

RNAP, PstI-digested template DNA and primer RNA were mixed in a molar ratio of 1:1.5:2 in the H buffer. RNAP was walked on the template DNA by adding the substrate ATP/UTP/CTP and GreA. The mixture was incubated for 20 minutes at 65°C, and loaded onto a Superose 6 gel filtration column (Cytiva). Fractions containing RNAP, DNA and full-length RNA were collected (Fig. S1B), concentrated and exchanged to Buffer C (20 mM Hepes-NaOH pH 7.5, 250 mM KCl, 5 mM MgCl_2_, 1% Glycerol), and protein amounts were estimated by SDS-PAGE. Approximately 20 pmol of EC was mixed with 25 pmol of Rho in the presence of 0.5 mM ADP·BeF_3_ in 400 μl of the buffer C. The sample was crosslinked with 3 mM of BS3 (Thermo Fisher) for 30 minutes at 30°C. The crosslinking was quenched by adding 100 mM of Tris-HCl pH 7.5. For the grid preparation, the samples were concentrated to about 20 ul. The samples were applied to Quantifoil R1.2/1.3 Copper 300 mesh grids (Quantifoil Micro Tools), and were plunge-frozen with a EM GP2 (Leica) at 10°C and 80% humidity.

### Cryo-EM Data collection and image analysis

Cryo-EM images were recorded by a Krios G4 transmission electron microscope (ThermoFisher)) equipped with BioQuantum energy filter and a K3 direct electron detector (Gatan). Two data set of ~13,000 and ~20,000 micrographs were collected (Table S1). Movies were aligned and dose-weighted using Relion3(*33*). CTF estimation was performed by gctf(*34*). The particle picking was performed with Warp(*35*) and Topaz(*36*), and the particle coordinates from each program were merged after 2D classification.

~1,641,000 particles of RNAP and RNAP-Rho complexes were selected by 2D classification (Fig. S1D), and subjected to 3D classification to discard particles without RNAP. Remaining ~997,000 particles were subjected to 3D refinement using an RNAP mask, followed by Bayesian polishing, local CTF estimation and beamtilt refinement to improve resolution. Then, particles from two data sets were merged for further 3D classification, which revealed classes of apo RNAP, EC and RNAP-Rho complexes 1 and 2 (Fig. S2). For this classification, a multi-template reference including type-1 and type-2 complexes, EC and apo RNAP templates, which was created using maps obtained from preliminary classifications, was utilized for better discrimination between the complexes.

For the type-1 complex, initial ~112,000 particles were refined with an overall mask, and then subtracted with a mask around Rho. Several rounds of 2D- and 3Dclassifications were performed to exclude particles with poor Rho density (Fig. S3A). ~43,000 particles with clear Rho density were selected for final reconstruction of a 4.0 Å map. As the quality of the overall reconstruction was compromised by the flexibility between Rho and the EC, independent reconstructions using tight masks around EC or Rho were also performed, respectively, and utilized for model building (Fig. S3A).

For the type-2 complex, ~269,000 particles from initial 3D classification were further classified using a mask around the complex 2 (Fig. S3B). A 5.8 Å map was reconstructed using ~147,000 particles.

~612,000 particles of EC and apo RNAP were re-classified through a 3D classification using a reference including EC and apo RNAP templates. ~220,000 EC particles and ~392,000 apo RNAP particles yielded 3.2 Å and 3.4 Å maps, respectively.

~693,000 particles of Rho-hexamer were selected by 2D classification (Fig. S1E), and subjected to 3D-classification (Fig. S2). ~239,000 particles were selected for 3D refinement with C6 symmetry enforced and subsequent Bayesian polishing. The particles were further 3D classified in C1 symmetry with symmetry relaxation by C6. ~205,000 particles were subjected to 3D refinement with C6 symmetry. Local CTF estimation and beamtilt refinement were performed to obtain a final 3.5 Å map.

### Model Building

The structure model of *T. thermophilus* Rho protomer was generated by homology modeling using *E. coli* Rho crystal structure16 (PDB 5JJI) as template and utilizing the SWISS-MODEL server(*37, 38*). The model of Rho-protomers excluding 59 residues at the N-terminal were fitted into the cryo-EM map of EC-unbound Rho and refined with Phenix real space refinement(*39*). For the model building of Rho-unbound EC and apo RNAP, the crystal structure of *T. thermophilus* EC (PDB 2O5I)(*24*) was fitted into the cryo-EM maps, and the models were refined with Phenix real space refinement. RNAP-Rho complexes models were generated by fitting the coordinates of EC-unbound Rho and Rho-unbound EC/RNAP into the maps. In the type-1 complex, structure models of RNAP-Rho interface (the β-flap tip, the β’ C-helix and ω C-helix of RNAP, and α18 helix of Rho protomer C) were built into the cryo-EM map using Coot and then refined with Phenix. All structure figures were created using ChimeraX.(*40*)

## Acknowledgments

The cryo-EM experiments were performed with the Krios G4 microscope at the RIKEN Yokohama cryo-EM facility.

## Author contributions

Conceptualization: YM, SS; Methodology: YM, HE, TY; Investigation: YM, MA, MG; Visualization: YM, SS; Supervision: SS; Writing—original draft: YM, SS; Writing— review & editing: YM, HE, SS

## Data and materials availability

The cryo-EM maps and coordinates have been deposited in the Electron Microscopy Data Bank (EMDB) and Protein Data Bank (PDB), respectively, with accession codes: XXXX, XXXX and XXXX.

