## Supplementary Materials for "Structural basis of the transcription termination factor Rho engagement with transcribing RNA polymerase"

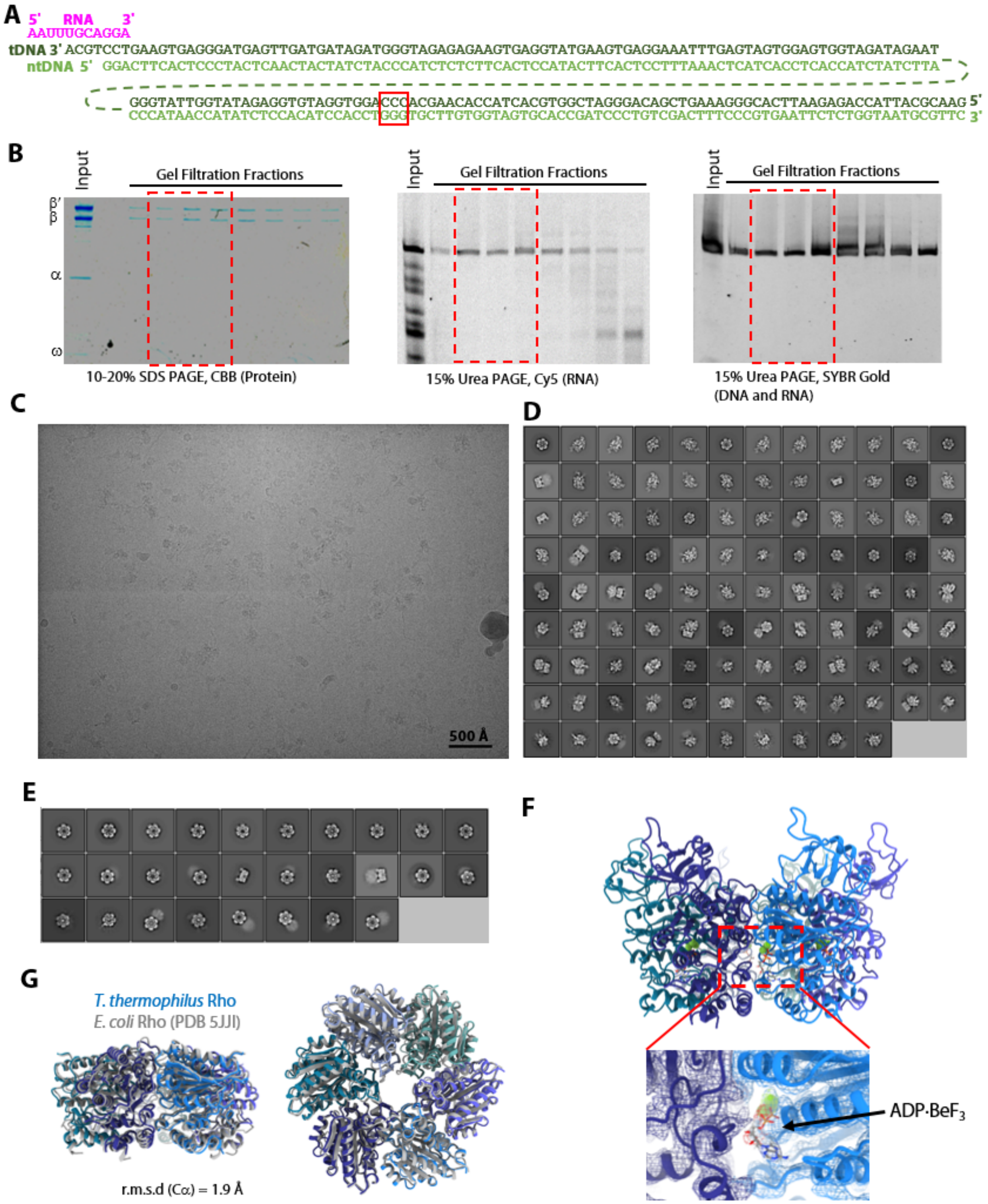


**Fig. S1. Sample preparation and image analyses of RNAP-Rho complex.**

(A) A schematic representation of the DNA and RNA for EC preparation. The G-triad is marked by a red box.

(B) Electrophoretic analysis of gel filtration fractions; proteins were detected with SDS-PAGE and DNA/RNA were detected with Urea-PAGE. Center and right are identical Urea-PAGE gels; after detection of fluorescent RNA (center), DNA and RNA were detected by staining gels with SYBR Gold (right). Fractions marked with red boxes were collected for cryo-EM sample preparation.

(C) A raw micrograph of a cryo-EM grid used for this study.

(D) 2D class averages for the RNAP and RNAP-Rho complex particles.

(E) 2D class averages for the Rho-hexamer particles.

(F) A close-up view of ADP·$BeF_3$ bound between Rho protomers A and B. The ADP is shown in stick model, and the beryllium and fluoride atoms are shown as spheres. The density map is shown as mesh colored to match the model.

(G) Superimposition of *T. thermophilus* Rho and *E. coli* Rho in the closed-ring form (PDB 5JJI). The models were superposed by their CTD region (Tth: residues 140-420, Eco: residues 131-413).

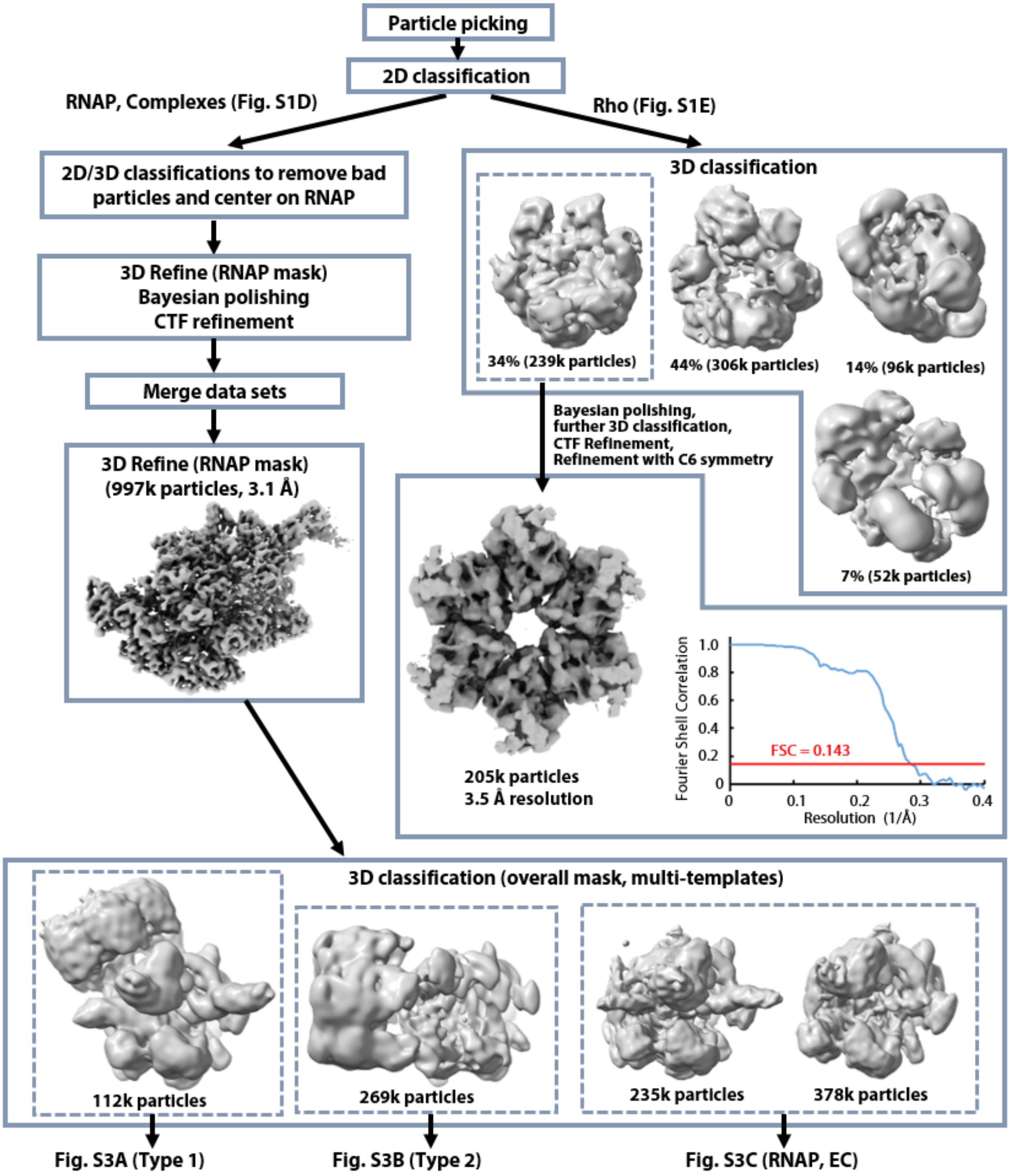


**Fig. S2. Workflow of the image analysis.**
Workflow of single-particle image analysis. Further analysis of particles containing RNAP are in Fig. S3.

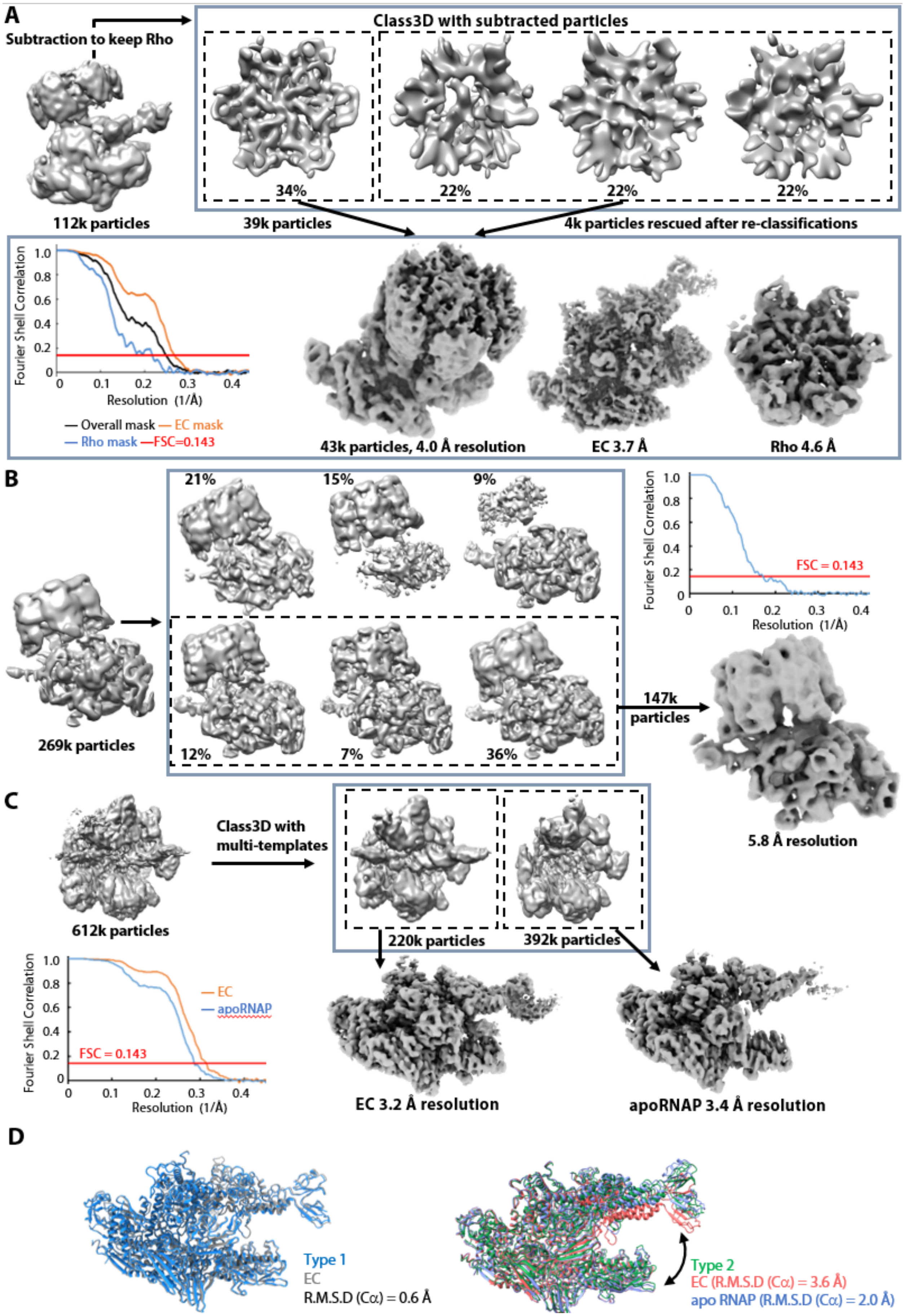
A
Subtraction to keep Rho
Class3D with subtracted particles
34%
22%
22%
22%
112k particles
39k particles
4k particles rescued after re-classifications
Fourier Shell Correlation
Resolution (1/Å)
— Overall mask — EC mask
— Rho mask —FSC=0.143
43k particles, 4.0 Å resolution
EC 3.7 Å
Rho 4.6 Å
B
21%
15%
9%
12%
7%
36%
269k particles
147k particles
FSC = 0.143
5.8 Å resolution
C
612k particles
Class3D with multi-templates
220k particles
392k particles
— EC
— apoRNAP
FSC = 0.143
EC 3.2 Å resolution
apoRNAP 3.4 Å resolution
D
Type 1
EC
R.M.S.D (Cα) = 0.6 Å
Type 2
EC (R.M.S.D (Cα) = 3.6 Å)
apo RNAP (R.M.S.D (Cα) = 2.0 Å)

**Fig. S3. Workflow of the image analysis and comparison of RNAP conformations in different complexes.**

(A) Image analysis of the type-1 complex.
(B) Image analysis of the type-2 complex.
(C) Image analysis of EC and apo RNAP.
(D) Comparison of RNAP conformation. Left: RNAP in the type-1 complex is superposed with that in EC by the RNAP core module (the two α subunits, residues 1-17, 394-700, 833-997 of the β subunit and residues 781-1069 of the β′ subunit). Right: RNAP in the type-2 complex is superposed with apo RNAP and EC by the RNAP core module.

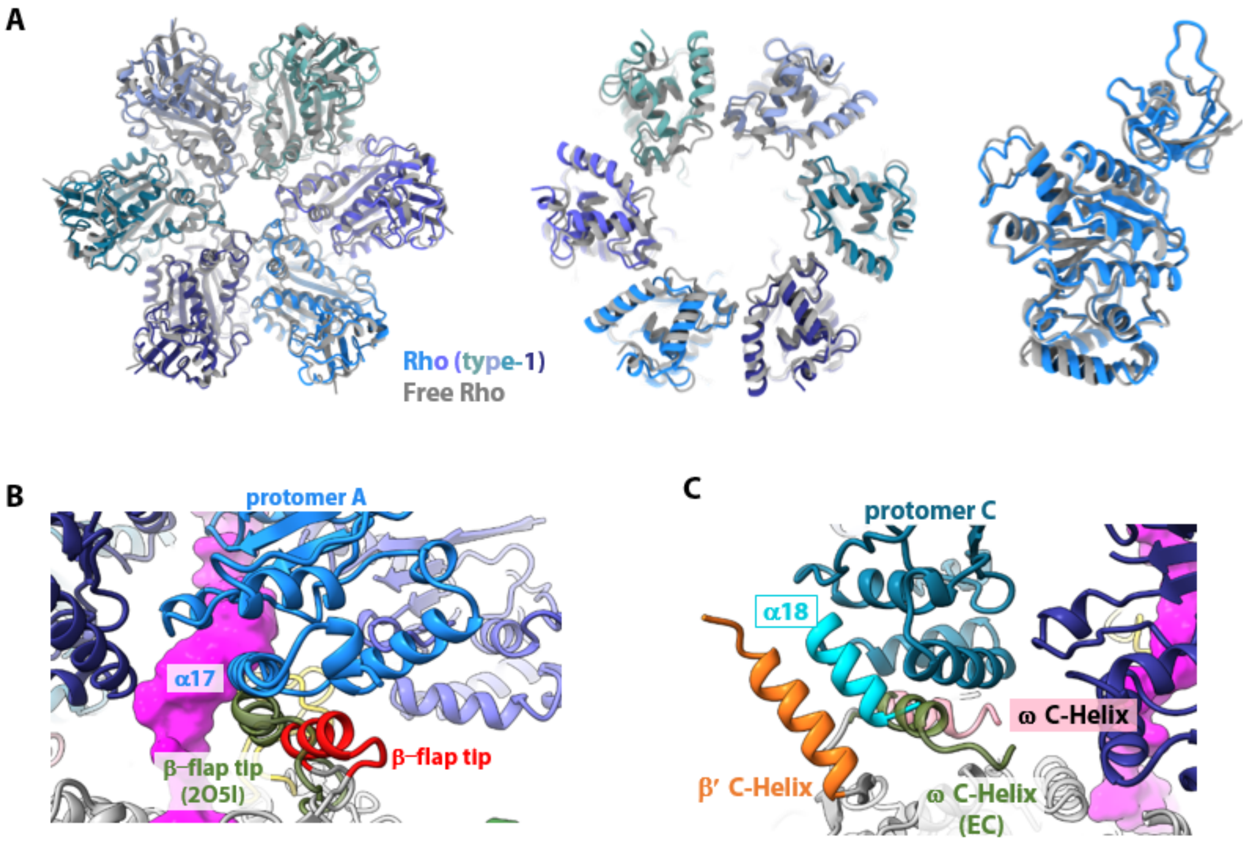


**Fig. S4. Structure of the type-1 complex.**

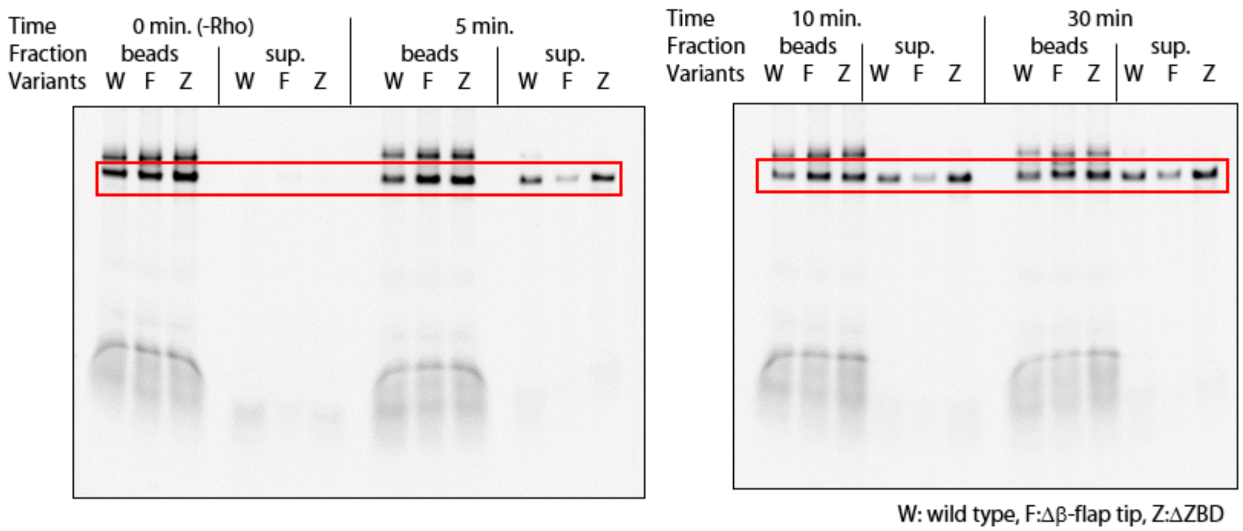


**Fig. S5. Gel images for the RNA release assay with RNA variants.**

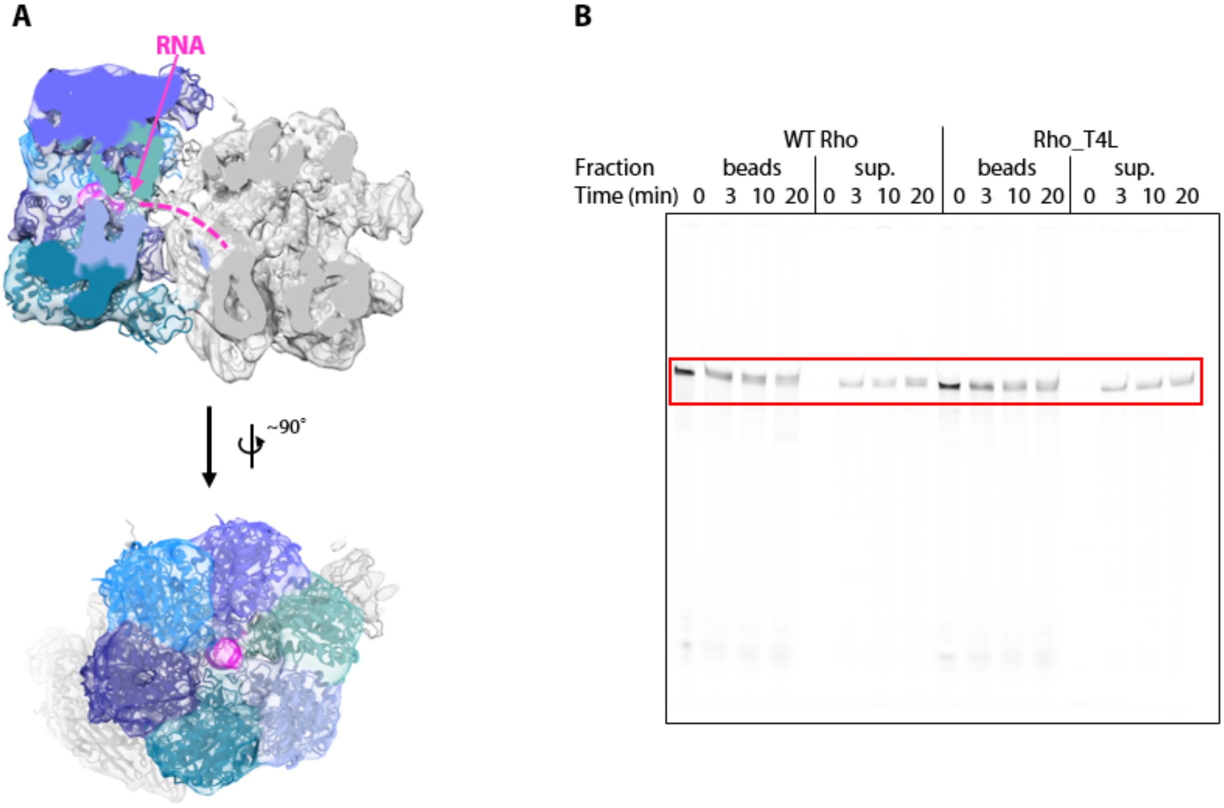


**Fig. S6. RNA density in the type-2 complex and a gel image of the RNA release assay with Rho_T4L.**

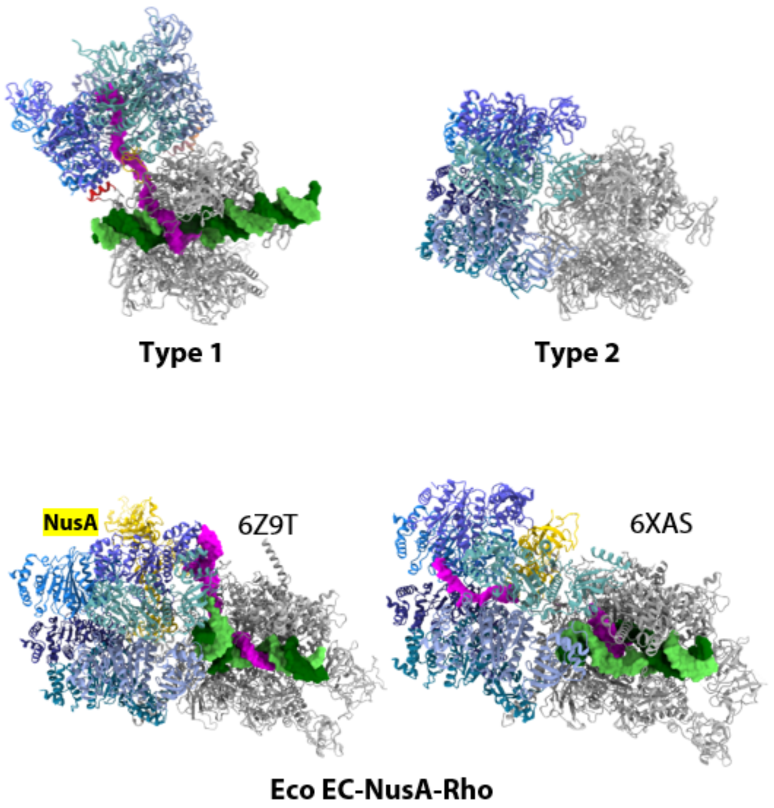


**Fig. S7. Comparison with *E. coli* EC-NusA-Rho complex structures.**

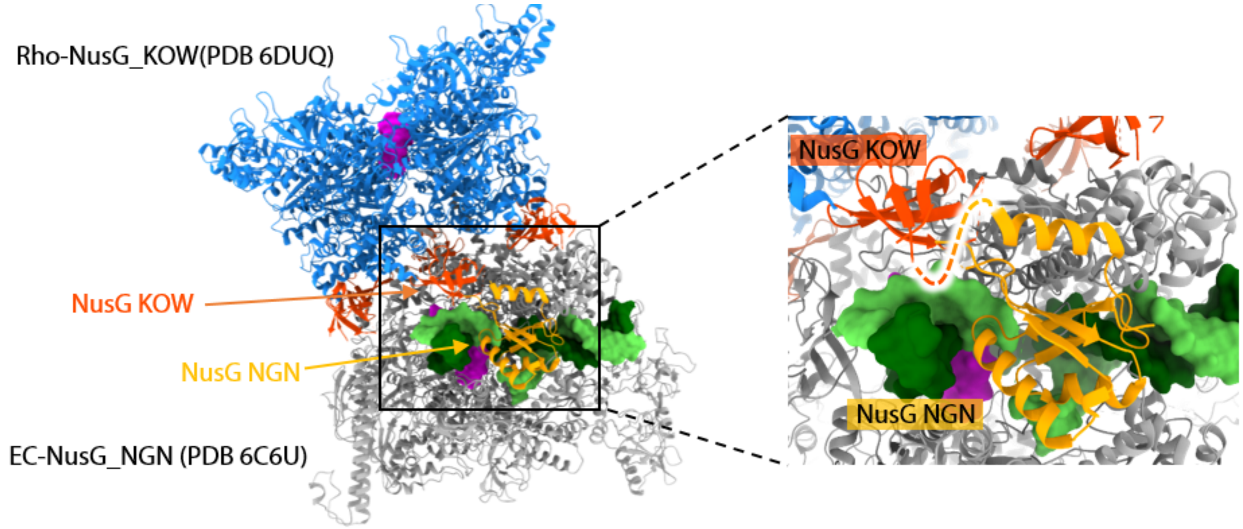


**Fig. S8. Modeling of NusG binding.**

The N-terminal (NGN) domain of NusG in the *E. coli* EC-NusG complex (PDB 6C6U)(*41*) was superimposed on the type-1 complex by the β′ coiled-coil (Tth: residues 539-583, Eco: residues 264-308). The C-terminal (KOW) domain of NusG in the *E. coli* Rho-NusG KOW domain complex (PDB 6DUQ)(*42*) was superimposed on the on the type-1 complex by CTD of Rho protomer F (Tth: residues 140-420, Eco: residues 131-413). The linker between the N- and C-terminal domains is depicted as a dotted line.

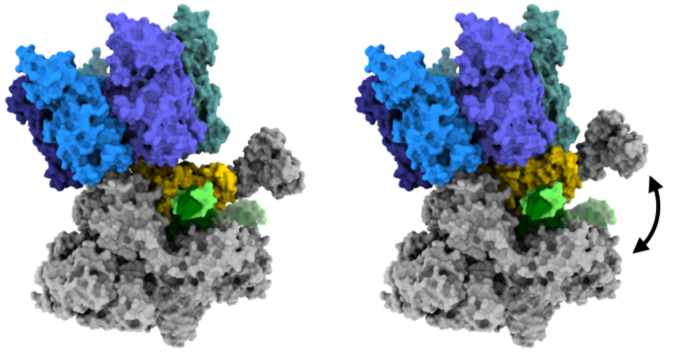


**Fig. S9. Modeling of clamp opening in the type-1 complex.**
Left: the type-1 complex; Right: apo RNAP was superimposed on the type-1 complex by the shelf module of RNAP (residues β 1006-1080, β′ 621-782 and β′ 1103-1430).

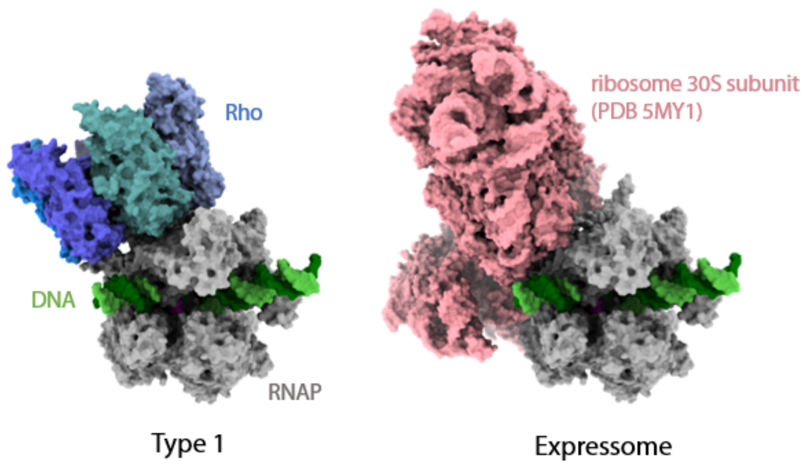


**Fig. S10. Overlapped binding sites of Rho and the ribosome on the EC.**
A model of the *T. thermophilus* “expressome” (right) was created by replacing *E. col*i RNAP in the *E. coli* expressome (PDB 5MY1)(*26*) by *T. thermophilus* RNAP.

### Table S1. Data Collection and image processing.

**Data collection and processing**

| | Dataset 1 | Dataset 2 |
|---|---|---|
| Number of micrographs | 13,764 | 19,776 |
| Magnification | 81,000 | |
| Voltage (kV) | 300 | |
| Pixel size (Å) | 1.06 | |
| Electron exposure (e–/Å$^2$) | 50 | |
| Defocus range (μm) | -0.9 ~ -2.7 | |

**Refinement**

| | Rho | EC | apo RNAP | Type 1 | | | Type 2 |
|---|---|---|---|---|---|---|---|
| No. particle images | 205,226 | 219,822 | 392,560 | 43,245 | | | 147,189 |
| | | | | composite | EC-mask | Rho-mask | |
| Map resolution* (Å) | 3.5 | 3.2 | 3.4 | 4.0 | 3.7 | 4.6 | 5.8 |
| CC mask | 0.65 | 0.78 | 0.79 | 0.60 | 0.82 | 0.62 | 0.58 |
| CC volume | 0.64 | 0.75 | 0.79 | 0.68 | 0.81 | 0.66 | 0.61 |
| Model composition | | | | | | | |
| Non-hydrogen atoms | 17,244 | 25248 | 23072 | 42819 | | | 40492 |
| Protein residues | 2,166 | 2971 | 2923 | 5143 | | | 5093 |
| DNA/RNA residues | 0 | 87 | 0 | 101 | | | 7 |
| Ligands | 18 | 3 | 3 | 21 | | | 21 |
| R.m.s. deviations | | | | | | | |
| Bond lengths (Å) | 0.004 | 0.003 | 0.004 | 0.005 | | | 0.007 |
| Bond angles (°) | 0.749 | 0.658 | 0.699 | 0.782 | | | 0.977 |
| Validation | | | | | | | |
| MolProbity score | 2.52 | 2.01 | 2.1 | 2.29 | | | 4.08 |
| Clashscore | 16.35 | 12.45 | 15.21 | 22.81 | | | 132.26 |
| Poor rotamers (%) | 4.93 | 0.04 | 0.08 | 0.02 | | | 17.18 |
| Ramachandran plot | | | | | | | |
| Favored (%) | 95.96 | 94.01 | 93.87 | 93.1 | | | 89.69 |
| Allowed (%) | 3.71 | 5.95 | 6.13 | 6.74 | | | 9.99 |
| Disallowed (%) | 0.32 | 0.03 | 0 | 0.16 | | | 0.32 |
| PDB ID | XXXX | XXXX | XXXX | XXXX | - | - | XXXX |
| EMDB ID | XXXX | XXXX | XXXX | XXXX | XXXX | XXXX | XXXX |

* FSC threshold = 0.143
